# Distinctive viral genome signatures are linked to repeated mammalian spillovers of H5N1 in North America

**DOI:** 10.64898/2025.12.09.693147

**Authors:** Matteo Chiara, Tommaso Alfonsi, Stefano Ceri, Erika Ferrandi, Anna Bernasconi

## Abstract

Highly pathogenic avian influenza H5N1 rarely infects mammals. In 2024–2025, however, genotypes B3.13 and D1.1 caused two independent spillovers into U.S. dairy cattle.

By analysing 26,930 complete H5N1 genomes from global surveillance, we identified 73 major viral groups, most of which show continent-specific distribution in Europe, Asia, Africa, and North America. North American viruses exhibit higher genetic diversity in specific viral segments, including variants potentially associated with mammalian adaptation. Both dairy-cattle-associated B3.13 and D1.1 genotypes originate from the same geographic macro-area, suggesting a possible regional hotspot where avian–mammalian interfaces may facilitate viral adaptation.

Our findings place the U.S. outbreaks in a global framework and indicate that North American H5N1 may be predisposed to cross-species transmission.

**Teaser:** Comparative genomics and geographic analyses delineate distinctive genomic features of H5N1 genotypes associated with U.S. dairy cattle spillover.

## Introduction

Influenza A viruses (IAVs) are among the most important zoonotic pathogens, causing recurrent epidemics and occasional pandemics in humans while also exerting a heavy toll on farmed animals’ health and food security (*1, 2*). Their segmented RNA genomes and broad host range facilitate repeated cross-species transmissions, making IAVs a constant concern for both veterinary and human medicine (*3, 4*). The highly pathogenic avian influenza H5N1 2.3.4.4b has circulated in wild birds and poultry for nearly three decades, causing repeated spillovers to mammals with sporadic but often severe outcomes (*5*). The capacity of H5N1 viruses to breach species barriers underscores the need for continuous genomic surveillance and careful analysis of viral evolution at a global scale (*6, 7*).

A compelling example of this risk emerged in the United States, where H5N1 has recently caused a large-scale epidemic in dairy cattle (*8*). Beginning in March 2024, an outbreak of respiratory illness and decreased milk production in dairy herds was traced to an H5N1 genotype designated B3.13 (*9*). This episode marked the first large-scale epidemic of influenza in cattle, raising urgent concerns about viral adaptation to mammals and the potential for onward spread and adaptation to humans (*10*). Comprehensive epidemiological and phylogenetic studies reconstructed the chain of B3.13 transmission and established that the outbreak most likely originated from a single spillover, with subsequent dissemination across state borders facilitated by the movement of animals and farm-related human activities (*9*). As the virus spread among cows, it acquired amino acid changes previously associated with mammalian adaptation (*11*).

In January 2025, D1.1 - a distinct H5N1 genotype-caused a second and independent spillover to dairy cattle and a second outbreak (*12*). Phylogenetic analysis showed that D1.1 and B3.13 are not related (*13*). Although smaller in scale compared with the B3.13 epidemic, the independent spillover of two distinct viruses to dairy cattle in a limited amount of time is unprecedented and raised significant concern (*13*).

These events raise several compelling questions. Why did dairy cattle in the United States become susceptible to two independent incursions of H5N1? Are the viruses circulating in North America more proficient at mammalian transmission? And can the geographic origin of these spillovers be pinpointed? Answering these questions is essential for assessing future risks of H5N1 adaptation to mammals and potential threats to public health.

So far, efforts to study the evolution and spread of H5N1 viruses associated with the recent dairy cow epidemic in the U.S. have relied primarily on viral genomes sampled in North America (*9*). These studies convincingly established the immediate progenitors of the cattle-adapted strains, but they left unaddressed the broader question of how these viruses relate to the global diversity of H5N1. Thousands of genomes have been sequenced across Europe, Asia, and Africa in recent years, yet these data have not been systematically compared with North American sequences.

Here, we address this gap by applying HaploCoV (*14*), a computational workflow for stratifying viral genomic diversity, to analyze H5N1 genomes worldwide. This approach delineates continent-specific viral groups beyond the current genotype-based nomenclature and traces the origin of both D1.1 and B3.13 to a shared area at the U.S.-Canada border.

Most strikingly, multiple internal segments forming the viral ribonucleoprotein (RNP) complex - PA, PB2, and NP - show increased diversity and distinctive genomic signatures in North America, distinguishing them from viruses elsewhere. Among these, nearly all North American strains carry a V105M amino acid substitution in the nucleoprotein gene. This change has been described in gain-of-function studies as one of a small set of mutations that facilitate influenza A virus adaptation to mammalian hosts (*15*). We show that it originated along the U.S. Atlantic coast in 2022 and subsequently became fixed in the North American viral population. The distinctive North American RNP complex gene repertoire warrants further study to clarify its origins and relevance for viral ecology and host range, with potential implications for risk assessment and surveillance.

## Results

### HaploCoV recapitulates and extends established genotype definitions

To establish a global framework for the comparative analysis of influenza A H5N1 diversity, we first applied HaploCoV to the complete dataset of 26,930 H5N1 genomes available in GISAID as of July 25th, 2025 (Data S1). HaploCoV stratifies sequence diversity by clustering viral genomes into groups based on pairwise genetic distances. The conceptual framework is illustrated in Fig. 1. First, the eight genomic segments were analyzed independently. Between six and fourteen groups were formed within every segment (Fig. 1A and Data S2), and large differences were observed in the numerosity of distinct groups (Fig. 1B). As exemplified in Fig. 1C, for every viral isolate, every segment was unambiguously assigned to one of the segment-specific groups. Concatenation of segment-level labels yielded genome-wide “segment-groups combinations”, hereafter referred to as HC comb. In total, 482 distinct HC comb were formed. Their frequency distribution was highly skewed: most of them (349) included at most 5 genomes, while a few of them (73), including 20 or more sequences, cumulatively accounted for 25,779 isolates (95.72% of the total). Only these 73 major groups were considered in subsequent analyses (Data S3). For these HC comb, the specific combination of segment-defined groups is represented by a distinctive color signature in Fig. 2A and reported in Data S4. As shown in Fig. 2B, most HC comb (63/73) were confined to a single continent, indicating limited evidence of intercontinental mixing. Only ten HC comb were detected across multiple continents (Fig. 2B), including three groups shared between Asia and North America, two groups shared between Europe and Africa, three distinct groups shared respectively between Asia and Europe, Europe and North America, and North and South America, and finally two distinct groups circulating across three continents: Asia, Europe and North America and Antarctica, North America and South America (see Fig. S1). These cross-continental distributions likely reflect sporadic, long-distance introductions mediated by migratory birds across both the East Atlantic and East Asia/Australasia flyways.

**Figure 1:**
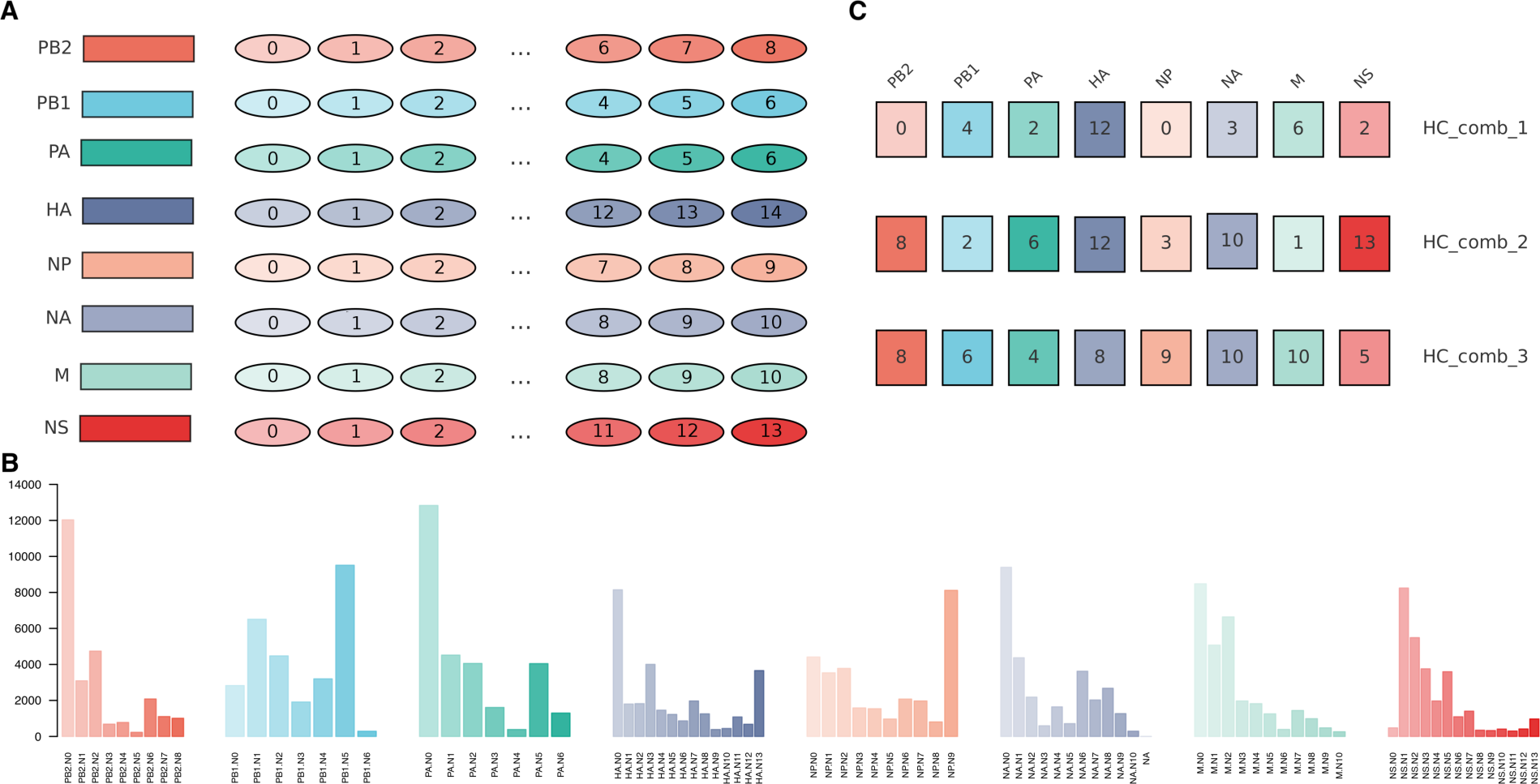
Application of HaploCoV to H5N1 clade 2.3.4.4b. (**A**) Conceptual workflow of HaploCoV applied to ↑26,800 viral isolates of H5N1 clade 2.3.4.4b collected worldwide between 2021 and July 25th, 2025, as available from the GISAID database. The eight genomic segments are represented as colored rectangles. For each segment, HaploCoV identified clusters of sequences, illustrated by ellipses, with the number of clusters varying by segment. (B) Bar plot showing the total number of sequences assigned to each HaploCoV-defined cluster across all segments. (C) Segment-level cluster labels were concatenated to generate a composite identifier (“HC comb”) for each virus. Only HC comb groups with more than 20 associated sequences/specimens were retained for downstream analyses.

**Figure 2:**
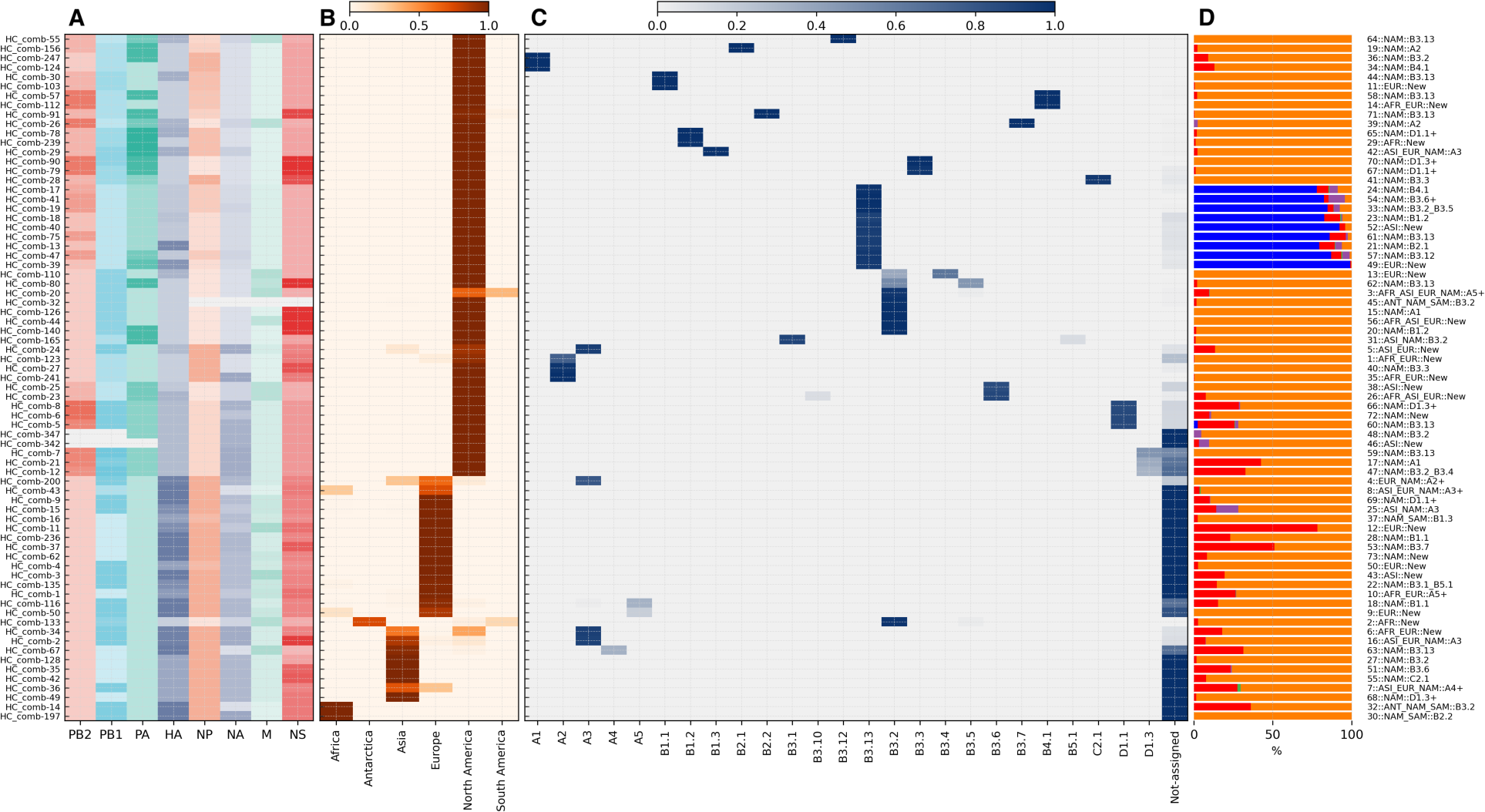
Salient features of HaploCoV-defined clusters in H5N1 clade 2.3.4.4b. (**A**) Heatmap representing the segment types across all eight genomic segments for each HC comb. Segment types are displayed using the same color scheme as in Fig. 1, with missing data indicated as blanks. HC comb identifiers are listed on the left. Complete data are provided in Data S4. (**B**) Heatmap showing the prevalence of each HC comb across continents, expressed as the proportion of total sequences per continent. Distinct associations between specific HC comb and continents are evident, with most detected in North America, followed by Europe and Asia. Some HC comb exhibit intercontinental circulation, such as HC comb-133 observed in both Antarctica and South America. (**C**) Heatmap illustrating the correspondence between HC comb and the established 2.3.4.4b genotype nomenclature. Values indicate the proportion (scaled to 1) of sequences in each HC comb assigned to a given genotype. “Not-assigned” denotes HC comb associated with sequences lacking genotype annotation. Most HC comb from North America correspond to a single genotype, while those from Europe, Africa, and Asia are frequently linked to new or unassigned groups. (**D**) Bar plot showing the proportion of host classes infected by each HC comb. Hosts are grouped into five broad categories: dairy cows, humans, other mammals, domestic birds, and wild birds. Several spillover events to poultry (red) are clearly visible in Europe and Asia. While spill-over to dairy cows (blue) is observed exclusively in North America. For each group, the corresponding names from the newly proposed system are indicated to the right of the bar plot.

Out of the 51 HC comb showing sustained circulation in North America, only three corresponded with more than one 2.3.4.4b genotype (Data S4), confirming that combinations largely recapitulate the previously established genotype-based nomenclature of 2.3.4.4b by Nguyen et al. (*9*) (Fig. 2C).

At the same time, HaploCoV increased resolution by partitioning some previously defined genotypes — including the cattle-associated B3.13 (nine groups) and D1.1 genotype (three groups) — into finer sub-groups, uncovering diversity not captured by the current 2.3.4.4b nomenclature. In total, 21 HC comb did not correspond to any defined H5N1 genotype (Fig. 2C and Data S4); these were observed in Europe (n=11), Asia (n=4), Africa (n=2) and North America (n=2), but also across continents (n=2) and represent a source of geographic and genomic diversity that was overlooked in previous studies.

Temporal stratification further underscored the distinct geographic patterns of 2.3.4.4b circulation across continents. By mapping the distribution of HC comb over time, we observed that different and unrelated groups dominated different continents in different years (Fig. S2). These dynamics emphasize that H5N1 circulation is not uniform worldwide but shaped by region-specific ecological and epidemiological processes.

Based on this evidence, we propose a scalable and rational set of rules for deriving informative group-level labels that can be integrated with the accepted genotype-based nomenclature of H5N1 2.3.4.4b:

1. First, each group should be assigned a progressive number. Numbering should follow chrono-logical order, with earlier groups receiving lower numbers;
2. Second, labels should incorporate geographic information indicating three-letter labels of the continent or continents in which widespread circulation was observed. Multiple continents should be separated by underscore (“_”);
3. Third, when a group corresponds to a previously defined genotype, this genotype should be explicitly included in the label. If no established genotype exists, the tag “New” should be used instead. When more than one genotype applies, they should be separated by underscore (“_”). Additionally, a + sign should be appended to the label if a group corresponding to a previously defined genotype contains 10% or more unassigned sequences.

For example, HC comb-9 is the sixth HC comb detected in chronological order, does not correspond to any previously defined genotype, and has been observed exclusively in Europe. According to our rules, it would be labeled: **6::EUR::New**. Complete data for all the viral isolates included in our analyses, along with labels compliant with the new proposed group-level labels, are reported in Data S3.

### Uncharacterized 2.3.4.4b viral groups from Asia, Europe, and Africa

A detailed analysis was performed on the 27 viral groups with sustained circulation in Europe, Asia, and Africa. Several of these 2.3.4.4b groups were associated with local spillovers to domestic birds and poultry (Fig. 2D and Data S5). In Asia (Fig. S3), we identified two previously undefined groups: 7::ASI::A4+, which circulated across multiple countries (China, Indonesia, Japan, Korea, Laos, and Vietnam) and was linked to recurrent poultry spillovers between 2021 and 2023 in Japan and Vietnam; and 38::ASI::New, which showed similarly widespread circulation and caused independent spillovers in comparable hosts and regions during the same period.

In Europe (Fig. S4), the autochthonous groups 6::EUR::New and 56::EUR::New have circulated across multiple European countries since 2021 and 2023, respectively, and continue to persist to the present. Group 6::EUR::New was responsible for major poultry outbreaks in the United Kingdom during 2021–2022 and for recurrent cases in the Czech Republic, whereas 56::EUR::New has been detected in poultry across several countries, including the Czech Republic, Italy, and the United Kingdom. Finally, we also observed group 49::EUR::New, which has been primarily associated with poultry outbreaks in the Czech Republic in 2024. Altogether, these groups represent regionally persistent 2.3.4.4b viral groups that have maintained sustained circulation within Europe.

In Africa (Fig. S5), we flagged 2::AFR::New, a group involved in poultry infections in Nigeria, Niger, and South Africa, with evidence of circulation from 2020 to 2024. These findings are consistent with recent epidemiological events (*16–18*) and underscore how HaploCoV-based stratification complements and extends existing surveillance systems by revealing unrecognized, region-specific emergent viral groups that warrant monitoring and formal classification.

### B3.13 and D1.1 origin in North America

We applied our high-resolution analytical framework to the 2.3.4.4b H5N1 clade to examine the genomic diversity and spatiotemporal dynamics of the B3.13 and D1.1 genotypes and their ancestors, B3.6 and A3. All the previously defined genotypes were subdivided into additional distinct groups; as shown in Fig. 3A and Fig. 3B, B3.6 was sub-divided into two groups, B3.13 into nine, A3 into four, and D1.1 into three groups.

**Figure 3:**
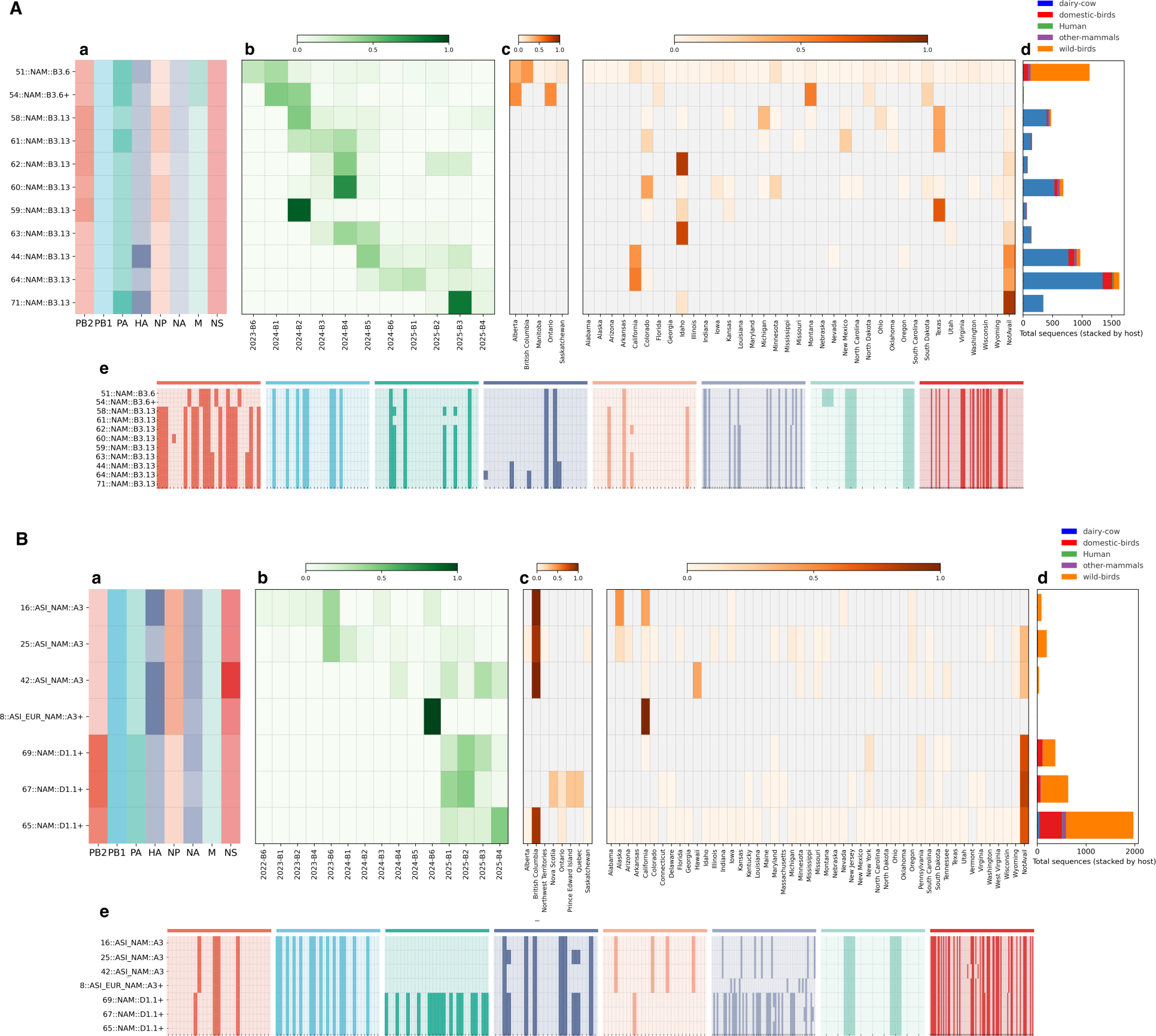
Circulation and genomic features of HaploCoV-defined groups in key 2.3.4.4b genotypes in North America. (**A**) HaploCoV groups corresponding to genotypes B3.6 and B3.13. Two distinct groups were resolved for B3.6. One group (54::NAM::B3.6+), detected only in Montana, Ontario, and Alberta, likely represents the progenitor of B3.13 based on shared HA segment composition. Eleven groups were identified within B3.13, many with strong geographic restriction; for example, 44::NAM::B3.13 and 64::NAM::B3.13 are specific to California. (**a**) Heatmap of segment composition, using the same color scheme as in Fig. 1. (**b**) Temporal circulation of each group, shown as the fraction of sequences per bimester; only bimester–group combinations with available data are displayed. (**c**) Geographic prevalence across Canada provinces and U.S. states, calculated as the proportion of sequences assigned to each group. (**d**) Bar plot showing the total number of sequences per group, stratified by host class: dairy cows, humans, other mammals, domestic birds, and wild birds. (**e**) Heatmap of non-synonymous substitutions across segments; dark shading marks substitutions, colored consistently with Fig.1. Distinct patterns across segments (e.g., in PB2 and NP between B3.6 and B3.13) are indicative of reassortment. Complete data are provided in Data S4 and Data S6. (**B**) Equivalent analyses for genotype A3 and D1.1. Four distinct groups were identified within A3; the new naming convention indicates their origin in Asia and subsequent introduction into North America. In D1.1, three distinct groups were detected, all restricted to North America. The 25::ASI NAM::A3 group is the most likely ancestor of D1.1 since it shares an identical HA segment. Sub-panels mirror those in (A). In panel (e), the distribution of non-synonymous substitutions is consistent with reassortment in PA, NP, and NA.

Groups defined within the B3.6 and B3.13 differed completely in the PB2 segment, and their NP segments also differed substantially, suggestive of reassortment of these two genome segments and consistent with phylogenetic analyses (*9*). Within B3.6, one group, 54::NAM::B3.6+, shared the same HA segment type as the early B3.13 groups (HA.N1, see Data S4). This pattern suggests that 54::NAM::B3.6+ probably represents the most likely ancestor of the B3.13 genotype. In B3.13, later groups, observed preferentially in California (44::NAM::B3.13 and 64::NAM::B3.13) and Idaho (63::NAM:B3.13), showed additional amino acid changes in the HA gene (discussed below). Unfortunately, although some of these groups are still circulating in 2025, their precise geographic distribution cannot be mapped due to missing metadata in the available records (see Data S4).

Across groups corresponding with the A3 and D1.1 genotypes, the PA, NP, and NA segments differed completely, indicative of reassortment and consistent with previous reports (*19*). Within A3, only one group, 25::ASI NAM::A3, shared the same HA segment subgroup as the D1.1 groups (HA.N3, Data S4), suggesting that this designation is the most plausible ancestor of the genotype for D1.1.

Interestingly, 25::ASI NAM::A3 the likely progenitor of D1.1 groups and 54::NAM::B3.6+, the ancestor of B3.13, were more frequently detected across a broad macro-geographic region along the eastern border between the United States and Canada - specifically, 25::ASI NAM::A3 in British Columbia and 54::NAM::B3.6+ in Montana (Fig. 5A-B). This pattern suggests that the ancestors of both D1.1 and B3.13 likely originated within a shared geographic zone spanning western Canada and the adjacent regions of the United States.

### Segment-level analyses reveal continent-specific diversity and mammalian-associated signatures

To complement the genome-level analyses, we examined each genomic segment individually to identify shared and genotype-specific patterns of diversity. The complete list of missense substitutions identified in every segment, and their distribution across distinct HaploCoV segment groups, is provided in Data S6. Overall, segment-level HaploCoV groups mirrored the geographic structure observed at the whole-genome level, with most diversity remaining locally constrained and distinct segment groups predominating in specific continents. In North America, we observed higher diversity in the internal genes NP, PA and PB2, and adaptive amino acid changes in viruses infecting dairy cattle in these segments and in the hemagglutinin (HA) gene. Here, we focus on these four segments, while results for the remaining genes are provided in the Supplementary Text and Figs. S6-S13.

### Hemagglutinin (HA)

HA sequences can be broadly divided into two major macro-groups based on their profiles of amino acid changes (Fig. 4A). The first comprises Eurasian groups (HA.N7, HA.N6, HA.N8, HA.N3, HA.N12, HA.N10, and HA.N13), with HA.N13 representing the earliest and most widespread group, detected across Africa, Asia, Europe, and the Americas. The second macro-group encompasses HA segments found predominantly in North America (HA.N11, HA.N2, HA.N9, HA.N0, HA.N5, HA.N1, and HA.N4). Integration of geographic and genomic data suggests that the HA groups predominant in North America most likely derive from HA.N0, which originated in Asia and reached the North America via Europe without establishing sustained circulation there. In addition, several independent introductions of Eurasian HA groups into North America—including HA.N13, HA.N12, and HA.N3 (the latter being observed only in America and Asia and associated exclusively with A3 and D1.1)—indicate that multiple entry events contributed to the observed North American HA diversity.

**Figure 4:**
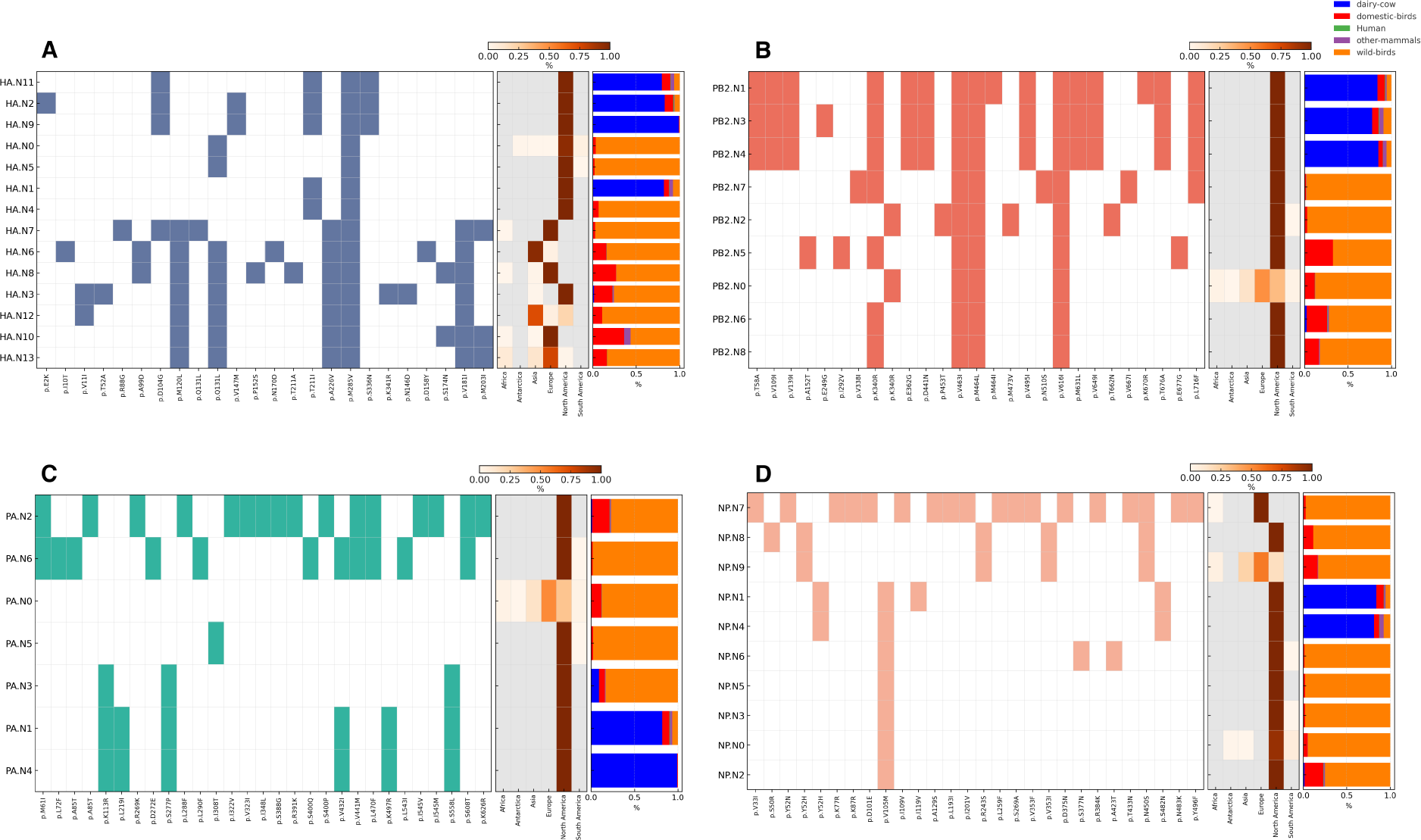
Characteristic substitutions, and host associations of segment groups in the HA, PB2, PA, and NP segments. For each segment, amino acid substitutions are shown as a heatmap. Substitutions are listed in the rows and reported in the format “reference residue–position–alternative residue.” White indicates the reference residue, while colored cells indicate the alternative residue. Segment-specific color schemes follow the conventions used in Fig. 1. Prevalence across continents and host range are shown as a heatmap and a bar plot, respectively, using the same formatting conventions as in Fig. 2. (**A**) HA: Eurasian macro-groups include HA.N13 (the earliest and most widespread), whereas specific North American groups (HA.N11, HA.N2, HA.N9) are linked to dairy cattle spillover and carry D104G and S336N, substitutions associated with mammalian virulence (see Fig. S6 for details). (**B**) PB2: two clusters are evident, one linked to the B3.13 cattle genotype (PB2.N1, PB2.N3, PB2.N4) and one including all the remaining PB2 segment-groups; most groups are observed exclusively in North America. All cattle-associated PB2s share M631L, enhancing polymerase activity in mammals (see Fig. S7). (**C**) PA: three groups can be cleary observed: PA.N2 and PA.N6; PA.N0 and PA.N5; PA.N3, PA.N1, and PA.N4. PA.N1 and PA.N4 (cattle-associated) carry K497R, which may act with PB2-M631L; PA.N2 is linked to D1.1/D1.3 poultry outbreaks (see Fig. S8). With the exclusion of PA.N5, all the groups are specific to North America (**D**) NP: three clusters: NP.N7; NP.N8 and NP.N0-NP.N6. NP.N0–NP.N6 all share V105M, and NP.N1/N4 carry Y52H, both linked to mammalian adaptation. While NP.N9 is more widespread worldide, the remaining NP groups are observed prevalently in North America (see Fig. S9).

Within North America, the HA groups associated with the dairy cattle spillover (B3.13 genotype) are particularly noteworthy, as they show the accumulation of several amino acid changes previously linked to mammalian adaptation. For instance, the HA groups HA.N11, HA.N2, and HA.N9—exclusively found in 44::NAM::B3.13, 64::NAM::B3.13, and 71::NAM::B3.13 (see Data S4), respectively—all carry the D104G substitution. This amino acid change has been reported to increase virulence in mammals (*20*) and is also present in HA.N7, an earlier and unrelated Eurasian group. The same HA groups also carry the S336N substitution, previously described as a low-frequency variant associated with increased virulence in mammals by Nguyen et al. (*9*). In addition to these group-specific changes, previous reports indicate that viruses linked with the dairy cattle outbreaks share two amino acid residues in the HA segment - 131Q and 211I - that could potentially expand the range of susceptible hosts. The 131Q residue - reported as Q131L in Fig. 4A because the reference genome used in our analysis already has 131Q-increases affinity for both α2,3- and α2,6-linked sialic acid receptors (*21*), whereas T211I broadens recognition of α2,3-linked glycans (*22*). We note that both 131Q and 211I also occur in HA.N4 and have already been identified in at least four distinct viral groups that predate the dairy cattle spillover (Data S4 and Data S6), indicating that these receptor-binding changes were established well before this event.

### PB2 and PA polymerase segments

The PB2 and PA segments display a skewed regional distribution (Fig. 4B-C), with the large majority of their HaploCoV-defined groups specific to North America (six of seven for PA, and eight of nine for PB2). In PB2, two main clusters can be distinguished based on shared patterns of amino acid changes: one associated with the B3.13 dairy cattle epidemic, characterized by multiple amino acid substitutions (PB2.N1, PB2.N3, PB2.N4), and another comprising all remaining groups. PB2.N0 is the only group detected across multiple continents—including Europe and Asia—and the most ancient in the dataset. Except for PB2.N2, which carries additional amino acid changes, all North American PB2 groups differ from PB2.N0 by a synonymous variant encoding the same residue (K340R).

All PB2 B3.13-associated groups carry the M631L substitution, which enhances polymerase activity in mammalian cells (*23*), but also several additional amino acid changes, which may warrant further functional investigation (T58A, V109I, V139I, E362G, D441N, V495I, V649I, and T676A). Concerning PA, three major PA groups can be distinguished. The first includes PA.N0, which is the most widespread and identical to the reference sequence used in this study, and PA.N5, which differs by a single amino acid change (I308T). The second group includes PA.N1, PA.N3, and PA.N4, which show a similar pattern of amino acid changes and are associated with the dairy cattle epidemic. Among these, PA.N3 is found in early B3.13 groups such as 61::NAM::B3.13, but also occurs in B3.6 (51::NAM::B3.6 and 54::NAM::B3.6+), whereas PA.N1 and PA.N4 are specific to B3.13 groups. Finally the last group includes PA.N2 and PA.N6, which are restricted to North America.

Both PA.N1 and PA.N4 carry K497R, which acts synergistically with PB2-M631L to fine-tune polymerase activity in mammalian hosts. This combination has been observed in several H5N1 isolates from mammals and is regarded as a potential molecular signature of mammalian adaptation (*23*). Additionally, we observed the V432I substitution, previously reported at low frequency but not functionally characterized (*9*).

The two remaining PA groups represent divergent groups observed exclusively in North America, possibly indicating the presence of a local viral reservoir. These include PA.N6 and, most notably, PA.N2, which is characteristic of D1.1, D1.3, and C2.1-associated groups.

### Nucleoprotein (NP)

Similar to PB2 and PA, the large majority of NP gene groups (8 out of 10) are restricted exclusively to North America (Fig. 4D). Patterns of shared amino acid substitutions delineate three main clusters. The first comprises NP.N8 and NP.N9: NP.N9 represents the earliest and most widespread group, whereas NP.N8 is confined to North America; the two differ by the single amino acid change S50R. The second cluster includes NP.N7, a divergent group observed exclusively in Europe. Finally, the third cluster is formed by groups NP.N0–NP.N6, which are all restricted to North America and share the defining substitution V105M. This amino acid change has been shown in gain-of-function studies to promote influenza A virus adaptation to mammals (*15*) and was already present in North American bird isolates by 2022, where it was first detected in the B1.1 2.3.4.4b genotype. Early occurrences were observed in viruses sampled in Florida and preceded by c.a. a month reports of IAV epidemic in dolphins (*24*). Within a few months, V105M rose to high prevalence and became fixed across all 2.3.4.4b NP genes in North America (Fig. 5C). The NP groups NP.N1 and NP.N4, associated with the B3.13 dairy cattle epidemic, both carry two additional amino acid changes, Y52H and S482N. The Y52H substitution enables the virus to escape recognition by mammalian restriction factors such as MxA and BTN3A3, thereby facilitating replication in mammalian hosts (*25*).

**Figure 5:**
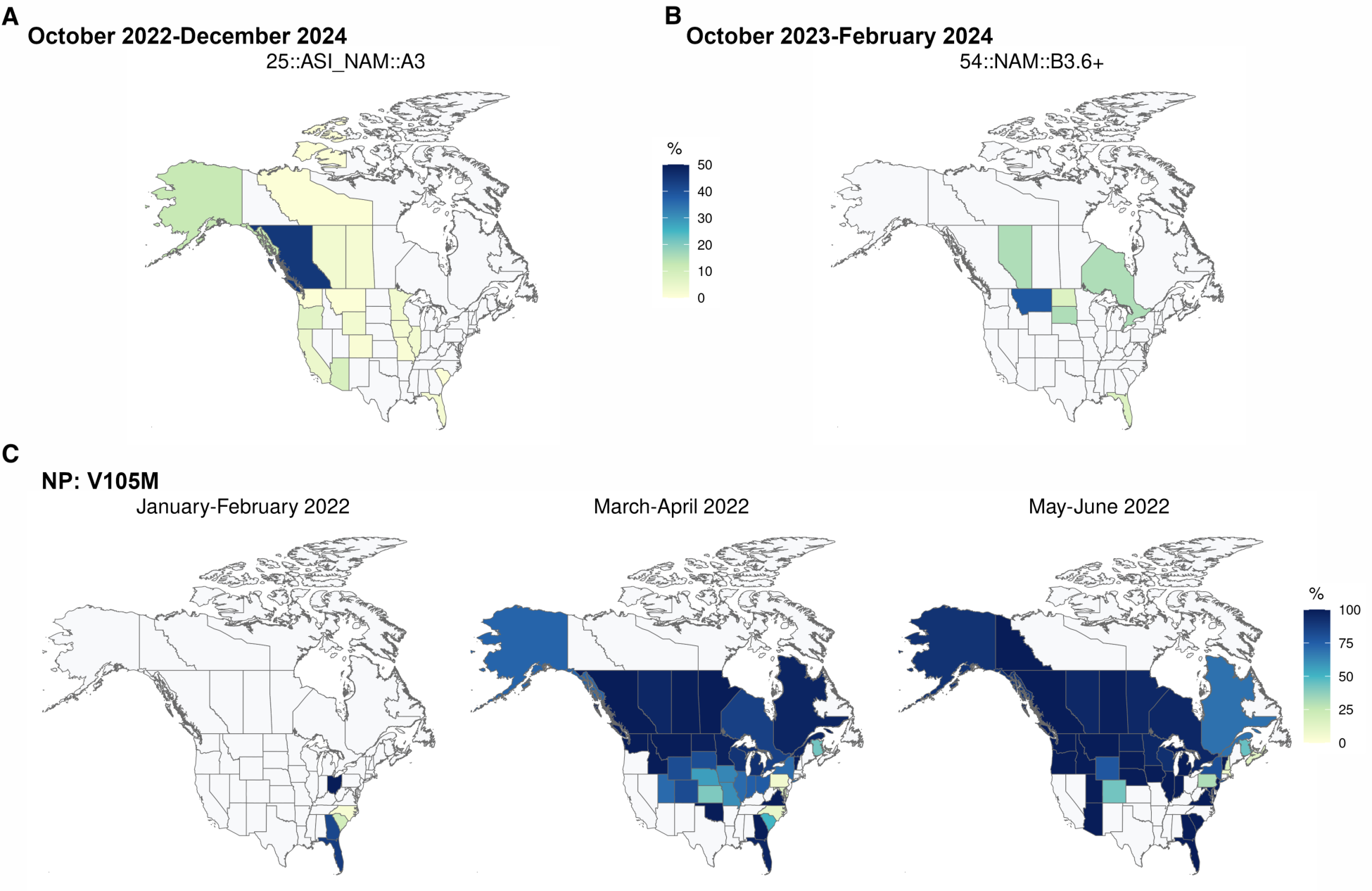
Geographic distribution of NP V105M and selected 2.3.4.4b viral groups in North America. (**A**) Map displaying the prevalence and geographic distribution of group 25::ASI NAM::A3, the likely progenitor of genotype D1.1, across Canada provinces and U.S. states from October 2022 to December 2024. Prevalence represents the percentage of sequences assigned to this group collected by state/province relative to the total number of isolates in the group. Values are capped at 50% to emphasize regions where the group was infrequently detected. (**B**) Equivalent prevalence map for group 54::NAM::B3.6+. This group and the one indicated in (A) circulated predominantly within a broad macro-geographic region spanning the northwestern border between the United States and Canada. (**C**) Maps showing the prevalence of the NP V105M amino acid substitution among North American H5N1 viruses of clade 2.3.4.4b. Prevalence is calculated relative to the total number of sequences collected in North America within each time interval shown and scaled by state/province. Three consecutive bimesters of 2022 are displayed, beginning from January 2022—the period when NP V105M was first detected. By March–April 2022, the majority of viruses collected in North America already carried this substitution, and its prevalence further increased in May–June 2022.

Taken together, these analyses highlight a consistent pattern across HA, PB2, PA, and NP segments, in which North American H5N1 viruses display increased diversity and share adaptive features that may enhance polymerase efficiency and host compatibility.

## Discussion

Our analyses provide a global perspective on the diversity and evolution of H5N1 viruses, situating the recent U.S. dairy cattle outbreaks within a broader comparative genomics framework. By applying HaploCoV to nearly 27,000 genomes, we confirmed that most viral diversity remains geographically constrained, but also identified rare intercontinental transfers that have repeatedly seeded North American 2.3.4.4b.

The resolution afforded by our study extends beyond conventional phylogenetic approaches by pinpointing the specific subgroups that gave rise to mammalian outbreaks. For B3.13, we identified 54::NAM::B3.6+ within the broader B3.6 genotype as the likely progenitor, based on shared HA segments and geographic circulation in several U.S. states and Canadian provinces prior to the cattle epidemic. This fine-grained resolution illustrates how local circulation of a relatively rare subgroup created the opportunity for spillover. For D1.1, although geographic metadata remains sparse, HaploCoV was able to isolate the subgroup responsible for the outbreak (68::NAM::D1.1+) and its most probable ancestor within the A3 genotype (25::ASI NAM::A3).

Together, these findings demonstrate the value of systematic, reproducible genotype classification in reconstructing the origins of reassortant genotypes and in identifying genetic backgrounds at heightened risk of cross-species transmission. Notably, the putative ancestral groups for both the B3.13 and D1.1 cattle-associated genotypes were detected within the same broad region of northeastern North America, indicating that this area constitutes a focal zone of viral circulation and diversification. Possibly due to ecological interfaces linking migratory wild birds, dense poultry production, and susceptible mammals in the Pacific flyway region encompassing British Columbia and the northwestern United States (*26, 27*).

Beyond reconstructing origins, our results highlight features of North American H5N1 viruses that may predispose them to mammalian adaptation. We show that North American viruses carry distinctive segment-level signatures, specifically in genes of the NRP complex, which may help explain why multiple independent spillovers into dairy cattle occurred within a single year. Although direct functional validation remains limited, the convergence of adaptive signatures across both surface and internal genes underscores the need for continued genomic and phenotypic surveillance of 2.3.4.4b viruses circulating in North America.

Furthermore, our framework extends genomic surveillance capacity globally by identifying viral groups not captured by the current genotype-based nomenclature but nonetheless associated with epidemiologically relevant events and independent spillovers to poultry in Asia, Africa, and Europe. These results demonstrate that HaploCoV-based stratification augments conventional surveillance approaches by identifying geographically restricted emergent groups that deserve continued surveillance and taxonomic recognition.

Our findings carry several implications. First, they underscore the importance of global data-driven surveillance: without comparing U.S. sequences against thousands of genomes from Eurasia and Africa, the unique features of North American viruses would have remained hidden. Second, they suggest that viral adaptation to mammals can occur in wildlife reservoirs before spillover is recognized, highlighting the need to monitor not only outbreak strains but also background genomic diversity in wild birds. Third, intercontinental geographic diversity suggests that interventions must be tailored by continent. While Eurasian and African viruses circulate extensively in poultry and wild birds, they have not (to date) acquired the NRP complex molecular signatures of viruses circulating in North America; close monitoring will be essential to avoid further spread.

There are, however, limitations. HaploCoV stratifies viruses based on sequence similarity and cannot itself demonstrate functional consequences of observed amino acid changes. Experimental studies are required to test whether North American 2.3.4.4b NRP genes are associated with enhanced replication or transmission in mammals. Metadata gaps also constrain interpretation: incomplete reporting of sampling locations, particularly in 2025, limits our ability to reconstruct precise geographic origins. Finally, surveillance remains uneven across continents, raising the possibility that additional introductions or adaptive changes have gone undetected.

In conclusion, our study shows that North American H5N1 viruses occupy a distinct evolutionary niche, marked by unique segment-level groups in NRP complex-associated genes and the fixation of a mammalian-adaptive NP amino acid change. These features likely contributed to the unprecedented emergence of influenza epidemics in dairy cattle. Ongoing surveillance and experimental validation will be critical to determine whether these findings represent an early warning of broader mammalian adaptation or whether they remain a regional anomaly. Either outcome underscores the urgency of sustained, globally coordinated influenza monitoring. These methods, however, should take into account the strong geographic differences in global viral circulation, local biases in data collection and curation, and should not rely solely on the existing nomenclature and lists of previously characterized amino acid changes. In this regard, focusing on the North American spillover, which has recently caused strong concerns (*9*), we recently developed a new computational method (*11*) capable of detecting the emergence and spread of novel viruses by analyzing exclusively compositional features of the viral genome, even at the level of individual sequences.

## Acknowledgments

We gratefully acknowledge all data contributors, i.e., the Authors and their Originating laboratories responsible for obtaining the specimens, and their Submitting laboratories for generating the genetic sequence and metadata and sharing via the GISAID Initiative, on which this research is based.

## Funding

The work was supported by Ministero dell’Università e della Ricerca (PRIN PNRR 2022 “SENSIBLE” project, n. P2022CNN2J), funded by the European Union, Next Generation EU, within PNRR M4.C2.1.1. Politecnico di Milano, CUP D53D23017400001; Università degli Studi di Milano, CUP G53D23006690001. AB Principal Investigator, MC co-Principal Investigator.

## Author contributions

M.C. data acquisition, data curation, data analysis, writing (first draft), funding acquisition; E.F. geographical data analysis, production of maps; S.C. research coordination; A.B. research coordination, funding acquisition. All authors jointly contributed to the interpretation, made a critical discussion of results, revised the draft, and approved the final version.

## Competing interests

The authors declare no competing interests.

## Data and materials availability

The findings of this study are based on metadata associated with 25,930 sequences available on GISAID up to July 25th, 2025, via https://doi.org/10.55876/gis8.251121ym. The code is provided on our Zenodo repository https://doi.org/10.5281/zenodo.17669841 (*28*).

## Supplementary materials

Materials and Methods

Supplementary Text

Figs. S1 to S13

Data S1 to S6

## Notes

https://doi.org/10.5281/zenodo.17669841

https://doi.org/10.55876/gis8.251121ym

### Competing Interest Statement

The authors have declared no competing interest.

