## SupplementaryMaterial for "Distinctive viral genome signatures are linked to repeated mammalian spillovers of H5N1 in North America"

**This PDF file includes:**

Materials and Methods

Supplementary Text

Figs. S1 to S13

Data S1 to S6 (captions)

### Materials and Methods

#### Data collection

We assembled a global dataset of complete H5N1 genomes ($n = 26{,}930$) deposited across 2021–2025 and for which at least five genomic segments (among PB2, PB1, PA, HA, NP, NA, M, NS) were available. Sequences and metadata (collection date, country, host) were retrieved from public repositories GISAID (*29*); data freeze: July 25th, 2025. Quality control retained sequences with segment length ≥95% of the expected reference and ≤1% ambiguous bases. *Host* was collapsed into five classes used throughout the study: dairy cattle, humans, other mammals, domestic birds, and wild birds.

#### HaploCov clusterization

Genomic diversity was stratified using *HaploCoV* (*14*), an unsupervised phenetic clustering framework for large-scale genomic surveillance of viral pathogens. The method identifies high-frequency non-synonymous variants from multiple sequence alignments and encodes their presence or absence across isolates to generate binary variant profiles. Phenetic clustering of these profiles defines "haplogroups"—sets of viral genomes sharing similar variant patterns—independent of existing nomenclature systems. *HaploCoV* can integrate with external classifications and annotate predicted functional effects of nucleotide substitutions.

To assess segment-level diversity, *HaploCoV* was applied separately to each genomic segment. Variants with a frequency greater than 0.01 were considered "frequent" and included in clustering analyses. Segment-specific groups were derived using parameters $N = 100$ and dist = 5, yielding distinct phenetic classes within each segment that capture patterns of recurrent variation, aiming to obtain a fine-scale discrimination of viral groups without forming an excessive number of groups. Reference sequences for all alignments and variant calling were obtained from the isolate *A/blue-winged teal/LA/22-033477-003/2022* (H5N1), corresponding to the genome assembly GCA_039319055.1 (GenBank). This isolate was also used as the reference in the *Flumina* workflow (*30*), and includes complete sequences for all eight segments (PB2, PB1, PA, HA, NP, NA, M, and NS). Segment-level alignments in this study were verified to be identical to those in the Flumina reference set.

For each isolate, segment-level group labels were reconciled to generate a composite identifier (`HC_comb`) representing the combination of segmental haplogroup assignments across the genome.

Only *major* `HC_comb` groups containing at least 20 sequences were retained for downstream analyses. This filtering yielded 73 `HC_comb` groups encompassing 95.7% of all genomes analyzed. These major groups were used for comparative analyses.

To evaluate concordance with established nomenclatures, each `HC_comb` was mapped to clade 2.3.4.4b genotypes (e.g., B3.13, D1.1, B3.6, A3) as defined by Nguyen *et al.* (*9*). For each `HC_comb`, the proportion of genomes assigned to each genotype was determined. Genotypes representing at least 10% of sequences within a given `HC_comb` were recorded; multiple genotypes were listed when more than one met this criterion. A complete mapping of `HC_comb` groups and corresponding genotypes is provided in Data S4.

#### Geographic and temporal analysis

Temporal prevalence was summarized in bimesters (two-month bins) to stabilize counts while preserving emergence dynamics. Geographic prevalence was computed as the proportion of sequences per group within each continent. For figures, we report: (i) heatmaps of defining non-synonymous substitutions (colored = present; white = absent), (ii) continental prevalence (row-normalized proportions), and (iii) host-class bar plots (five host classes). Cross-continental occurrences were flagged when the same group was observed in ≥2 continents with a prevalence above 0.1. Because sampling intensity differs across regions and hosts, analyses emphasized qualitative patterns—such as emergence, restriction to North America, or repeated introductions—rather than formal statistical testing in the main text.

All visualizations were generated in Python using `matplotlib`, with color palettes matching the `ggsci` package (R). Temporal data were handled with the `lubridate` R package to standardize and aggregate collection dates.

Choropleth maps and barplots summarising the geospatial distribution of selected groups/amino acid changes in time periods of interest were generated in R using the `ggplot2` package. Reference data for worldwide political administrative boundaries were retrieved from the `geoBoundaries` database (*31*) through the dedicated R client (`rgeoboundaries`).

#### Segment-specific analysis

For each segment, we enumerated the characteristic amino acid substitutions per group, as identified by HaploCoV and summarized: (i) emergence time (first detection by bimester), (ii) continental distribution, and (iii) host associations. Amino acid positions are reported relative to the reference isolate *A/blue-winged teal/LA/22-033477-003/2022 (H5N1)* (GCA_039319055.1), unless otherwise noted.

### Supplementary Text

#### Detailed geographic distribution and prevalence of HaploCoV segment groups for NS, M, PB1, and NA segments

**Non-structural protein (NS).** Unlike other segments, the NS gene shows limited geographic bias and no specific associations with host shifts or epidemiologically relevant events, suggesting a largely conserved role of NS in maintaining basic viral functions rather than driving adaptation. Four main groups can be identified based on patterns of amino acid changes (see Fig. S10). NS.N6 is the earliest detected segment and shares multiple substitutions with NS.N12, NS.N3, NS.N5, NS.N0, NS.N10, and NS.N2. With the exception of NS.N10 and NS.N3, which are specific to North America, most designations in this group show a broad geographic distribution and are observed across two or more continents. The second cluster includes NS.N8, NS.N7, NS.N11, NS.N1, and NS.N13. Clusters 1 and 2 share 18 defining residues, and both display widespread circulation, indicating a globally conserved evolutionary trajectory for this segment. The third cluster comprises only NS.N9, which differs by two amino acid changes compared with the reference sequence and is observed primarily in Europe and Asia. The fourth and last cluster is represented by the divergent NS.N4, observed exclusively in Europe.

**Matrix protein (M).** Two main clusters can be defined in the M segment based on shared amino acid changes (see Fig. S11). The first includes M.N2, M.N9, M.N0, M.N6, M.N3, M.N10, M.N4, and M.N5, all of which share the S85N substitution and exhibit a widespread distribution across different continents. M.N0 represents the earliest and most broadly distributed group. The second cluster includes M.N8, M.N1, and M.N7, which share the N87T substitution, while M.N1 and M.N7 carry an additional A227T change. Among these, M.N1 is specifically associated with the B3.13 genotype and the dairy cattle epidemic. Although neither N87T nor A227T has been functionally characterized in H5N1, their recurrent appearance in North American viruses and association with the dairy cattle viruses suggest that these substitutions may represent early steps in local diversification. Nevertheless, they should be considered candidate adaptation markers rather than established mammalian-adaptive signatures.

**Polymerase basic 1 (PB1).** In PB1, two main clusters of segment groups can be distinguished based on shared profiles of amino acid changes (see Fig. S12). The first includes PB1.N2, PB1.N0, PB1.N5, and PB1.N6, which share seven defining amino acid changes and are globally distributed across two or more continents—with the exception of PB1.N6, which is specific to North America. The second cluster comprises PB1.N1, PB1.N4, and PB1.N3, all defined by the D264E substitution. PB1.N1, which is associated with the B3.13 genotype, displays several additional amino acid changes compared with the other PB1 groups. These include E75D, M171V, and A587P, none of which have been experimentally characterized. Current polymerase studies and PB1 mutational constraint maps do not flag these residues as known adaptation sites, and they should therefore be regarded as candidate variants for future investigation. Taken together, the PB1 diversity observed here mirrors the patterns seen in PB2 and PA, with North American viruses showing unique combinations of substitutions that may influence polymerase activity and replication efficiency in mammalian hosts.

**Neuramidase (NA).** Observed patterns of nonsynonymous defining substitutions in the NA segment revealed four major macro-groups, each characterized by shared sets of amino acid substitutions (see Fig. S13).

Within these, the group formed by NA.N5 and NA.N8 emerged as uniquely North American. NA.N5 and NA.N8 exhibited a higher proportion of nonsynonymous substitutions than other clusters; notably, NA.N8 was associated with the D1.1 genotype responsible for the dairy cattle spillover in January 2025 (*19*). Both NA.N5 and NA.N8 appeared only recently in the temporal record (see Fig. S13). Although the functional impact of their changes remains to be determined, these clusters represent strong candidates for further investigation.

Another group, included NA.N1, NA.N3, NA.N2 NA.N0, NA.N10, NA.N6 and NA.N7, which all share the A52T and L399W amino acid changes. While NA.N0, NA.N7 and NA.N6 show a widespread geographic distribution and were detected across most continents, NA.N1, NA.N2 and NA.N3 are observed almost exclusively in North America. NA.N1 and NA.N2 are both linked to the B3.13 genotype and share several private amino acid variants (L269M, V321I, and S339P). In addition, NA.N1 carries the N71S substitution, located near the enzymatic head domain and previously associated with subtle alterations in neuraminidase stability and antigenicity. Interestingly, the majority of North American NA groups also share the nonsynonymous T405S substitution within the second sialic acid–binding site (2SBS), a region that modulates the functional balance between HA and NA during receptor binding. While the precise role of T405S remains unknown, substitutions in this domain have been linked to host-range adaptation, warranting further study.

The remaining two groups are formed by NA.N4, which is observed prevalently in Europe but also in Asia, and by NA.N9 that crossed the inter-continental boundaries of North America, Asia and Europe. Both groups carry defining aminoacid substitutions that are not observed elsewhere.

Collectively, these observations suggest that NA diversification is largely structured by geography, with each continent maintaining its own prevalent variants. The North American-specific NA clusters might represent either independent introductions of unsampled Eurasian segment-groups and/or local NA segment variants that are not observed outside of North America. Since the origin of D1.1 is clearly in North America, we believe that at least for NA.N8, the latter represents the most likely scenario.

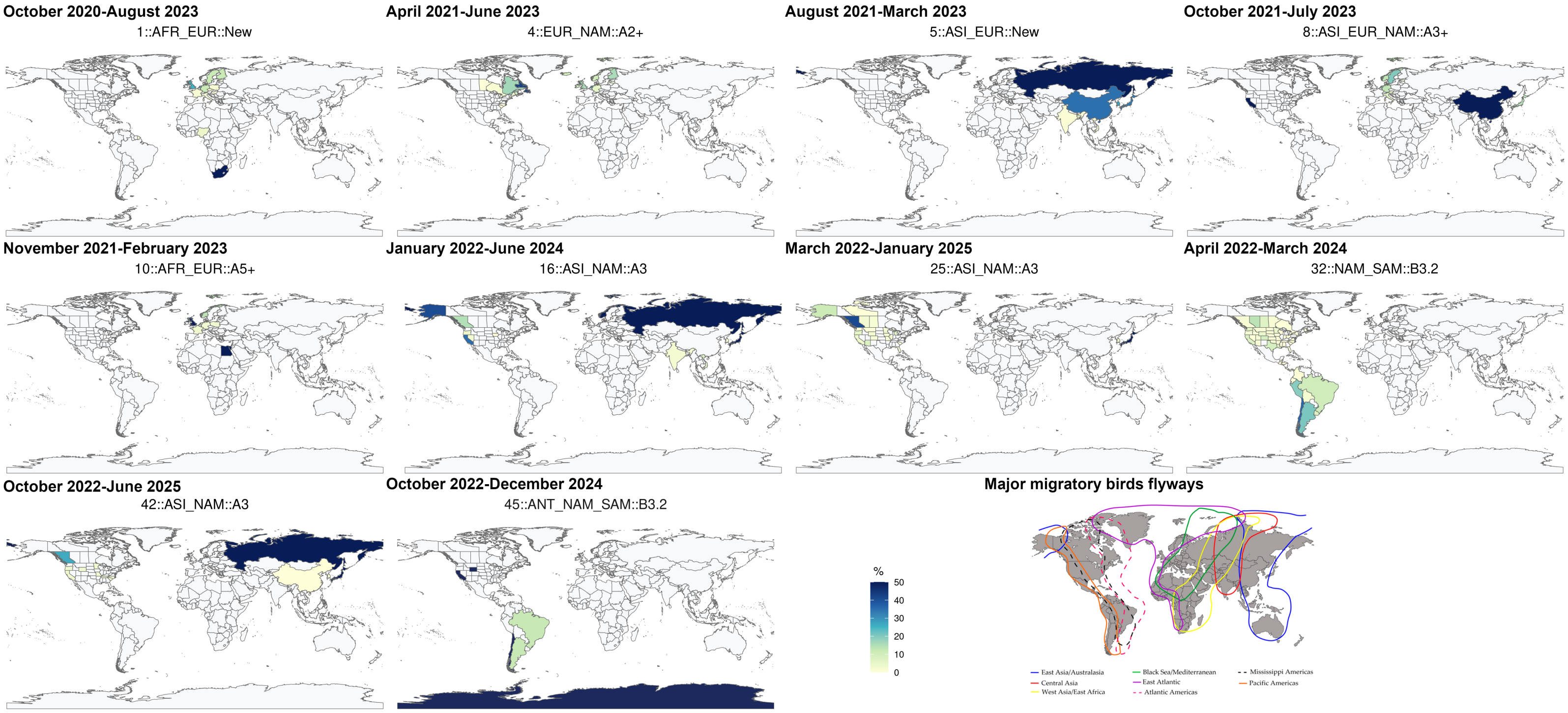


**Figure S1**: **Prevalence and geographic distribution of viral groups across continents.** For each of the ten viral groups with intercontinental circulation, maps show the prevalence of the group across regions or states where specimens were collected. Prevalence is computed with respect to the total number of sequences assigned to each group, and for every state or region is scaled relative to the corresponding continent. Values are capped at 50% to highlight areas where the group was infrequently detected. A map of major bird migratory flyways is also added as a reference for comparison. This map was obtained and adapted from (*32*).

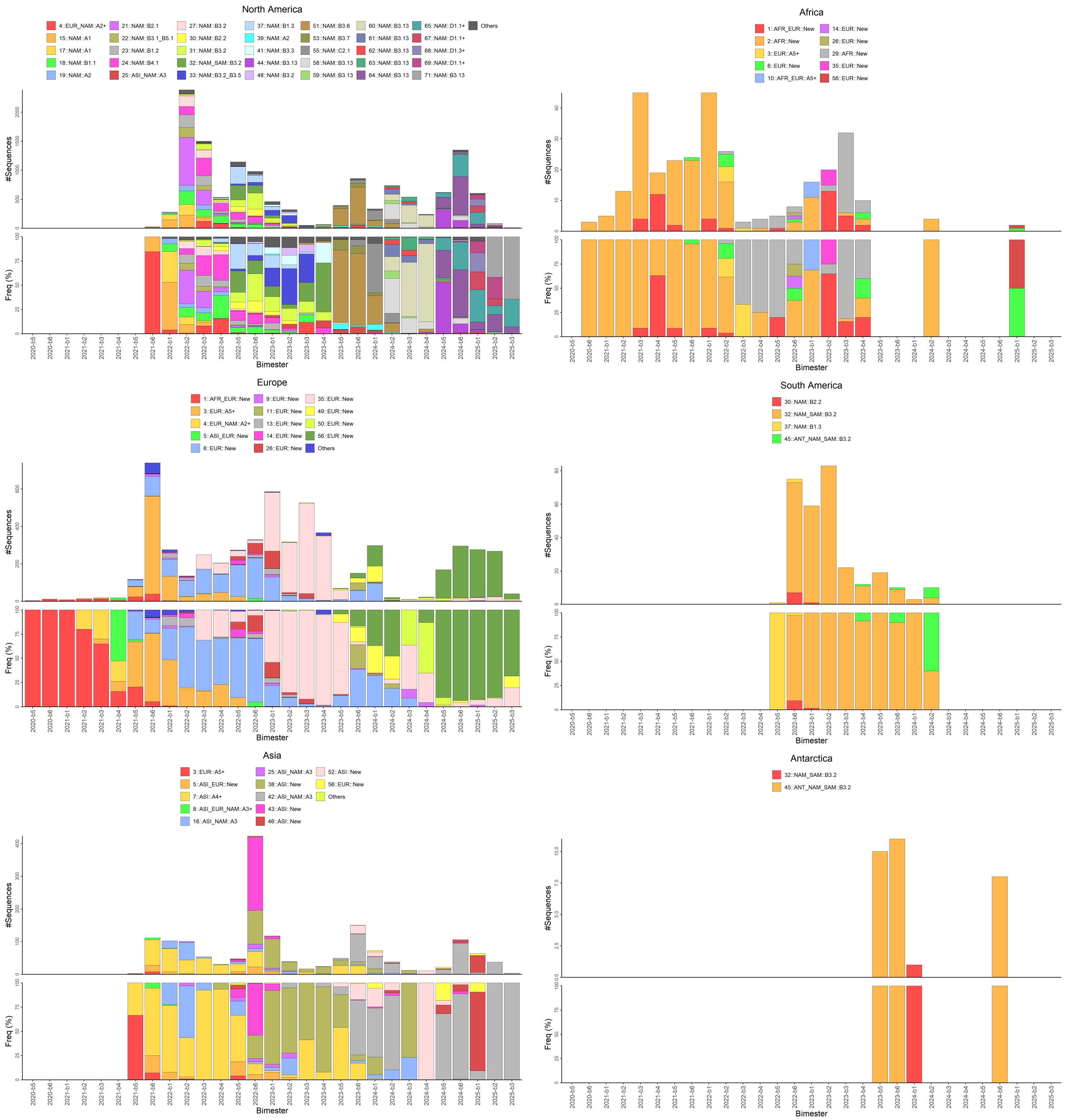


**Figure S2**: **Temporal prevalence of viral groups across continents.** Bar plots display the number and prevalence of viral groups across continents and bimesters. Prevalence is calculated relative to the total number of sequences available within each bimester and is scaled by continent. Only groups reaching a minimum prevalence of 5% in at least one of the analyzed time periods are shown explicitly; all remaining groups are aggregated into the “Others” category.

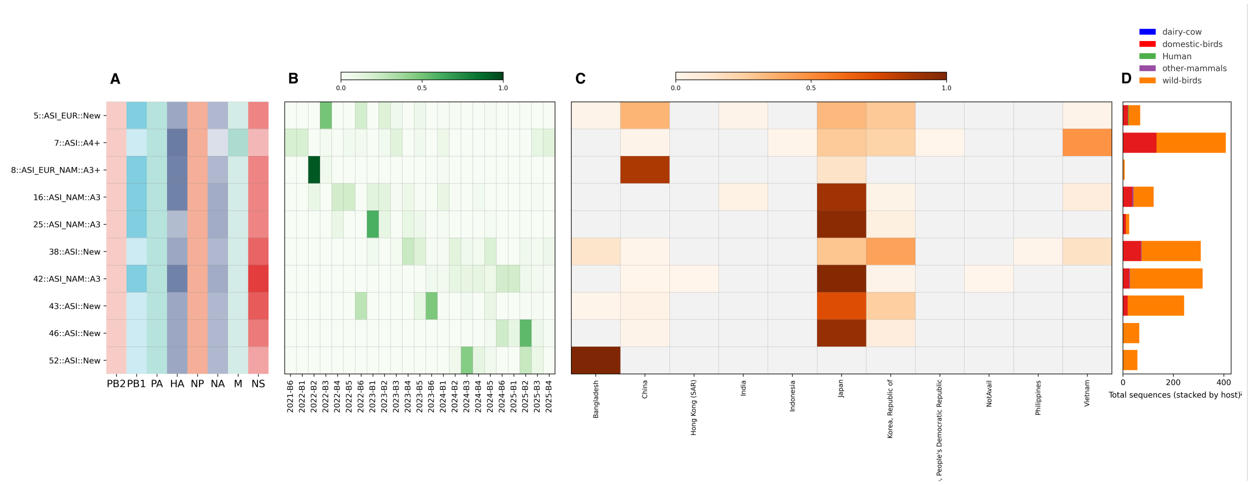


**Figure S3**: **Genomic diversity of viral groups in Asia.** (**A**) Heatmap showing segment types across all eight genomic segments, colored as in Fig. 1. Group names follow the newly proposed nomenclature (see Data S4). (**B**) Temporal circulation of each group, expressed as the fraction of sequences per bimester; only bimester–group combinations with available data are shown. (**C**) Geographic distribution of each group across countries where at least one sequence was detected; gray indicates absence. (**D**) Bar plot showing the total number of sequences per group, stratified by host class (dairy cattle, humans, other mammals, domestic birds, and wild birds).

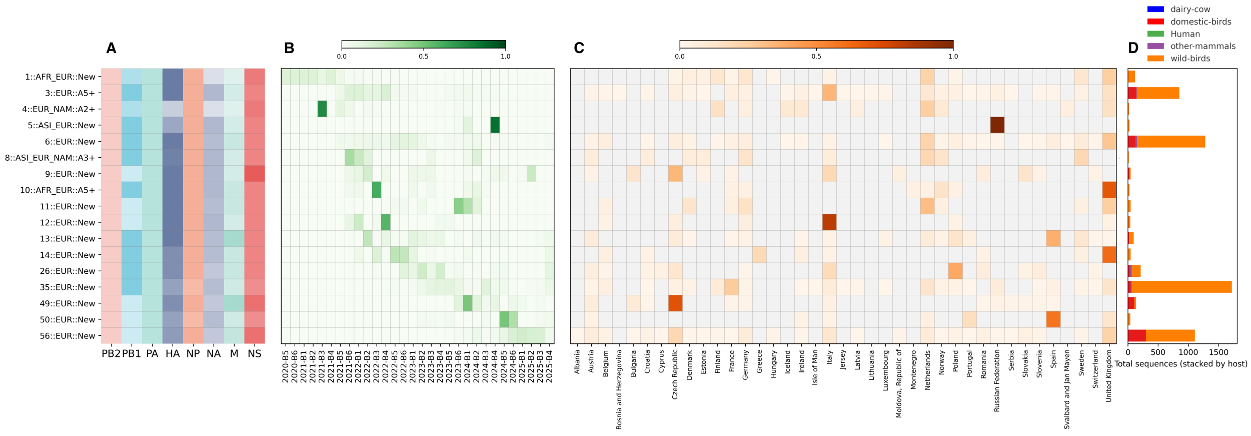


**Figure S4**: **Genomic diversity of viral groups in Europe.** (**A**) Heatmap showing segment types across all eight genomic segments, colored as in Fig. 1. Group names follow the newly proposed nomenclature (see Data S4). (**B**) Temporal circulation of each group, expressed as the fraction of sequences per bimester; only bimester–group combinations with available data are shown. (**C**) Geographic distribution of each group across countries where at least one sequence was detected; gray indicates absence. (**D**) Bar plot showing the total number of sequences per group, stratified by host class (dairy cattle, humans, other mammals, domestic birds, and wild birds).

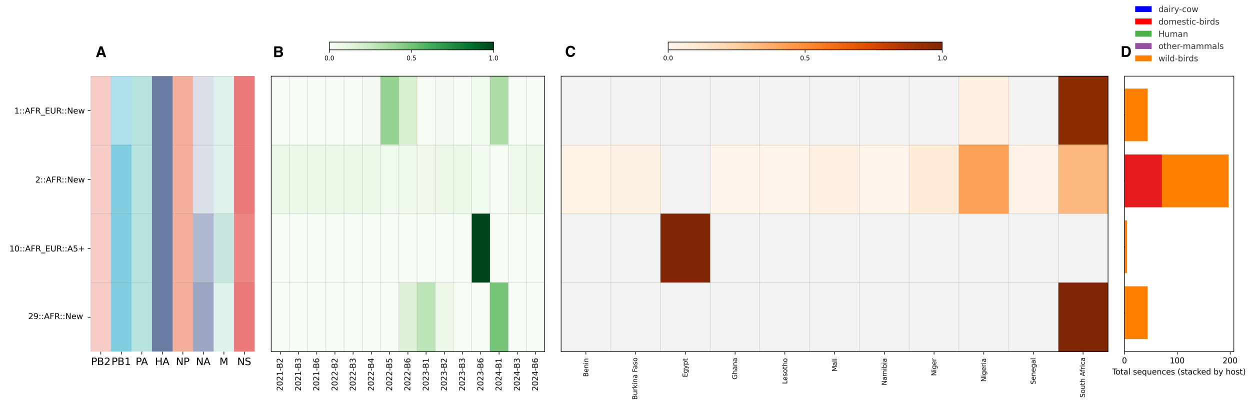


**Figure S5**: **Genomic diversity of viral groups in Africa.** (**A**) Heatmap showing segment types across all eight genomic segments, colored as in Fig. 1. Group names follow the newly proposed nomenclature (see Data S4). (**B**) Temporal circulation of each group, expressed as the fraction of sequences per bimester; only bimester–group combinations with available data are shown. (**C**) Geographic distribution of each group across countries where at least one sequence was detected; gray indicates absence. (**D**) Bar plot showing the total number of sequences per group, stratified by host class (dairy cattle, humans, other mammals, domestic birds, and wild birds).

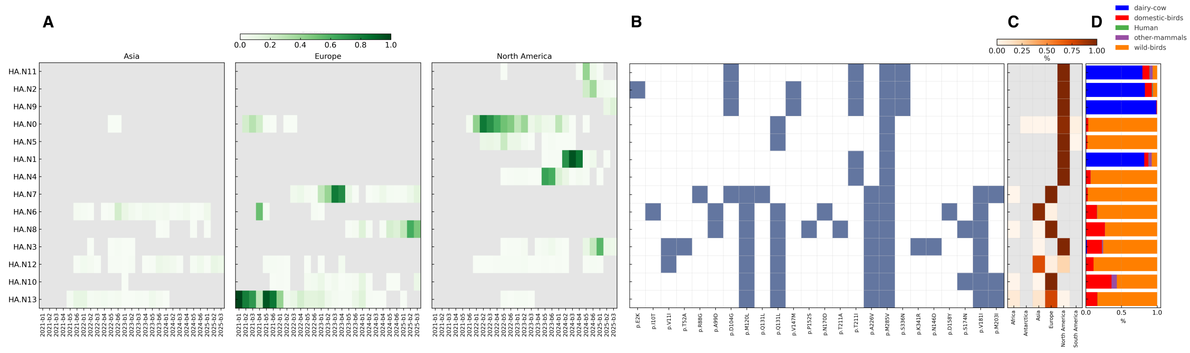


**Figure S6**: **Hemagglutinin (HA): temporal, geographic, and host distribution of groups.** (**A**) Temporal circulation of HA groups, expressed as the fraction of sequences per bimester. (**B**) Heatmap of characteristic non-synonymous substitutions showing two major macro-groups: North American (HA.N11, HA.N2, HA.N9, HA.N0, HA.N5, HA.N1, HA.N4) and Eurasian (HA.N7, HA.N6, HA.N8, HA.N3, HA.N12, HA.N10, HA.N13). (**C**) Geographic prevalence highlights multiple introductions from Eurasia into North America, including HA.N13, HA.N12, and HA.N3. (**D**) Host association barplot showing three dairy cattle–linked groups (HA.N11, HA.N9, HA.N2) carrying D104G and S336N; L131Q and T211I, which may expand receptor binding, were already present in HA.N4 (B3.6/B3.7). Note: In the text, the substitution is described as L131Q, while in the heatmap it appears as Q131L because Q131 is fixed in the reference sequence used for the analysis; amino acid changes are therefore reported relative to this updated reference.

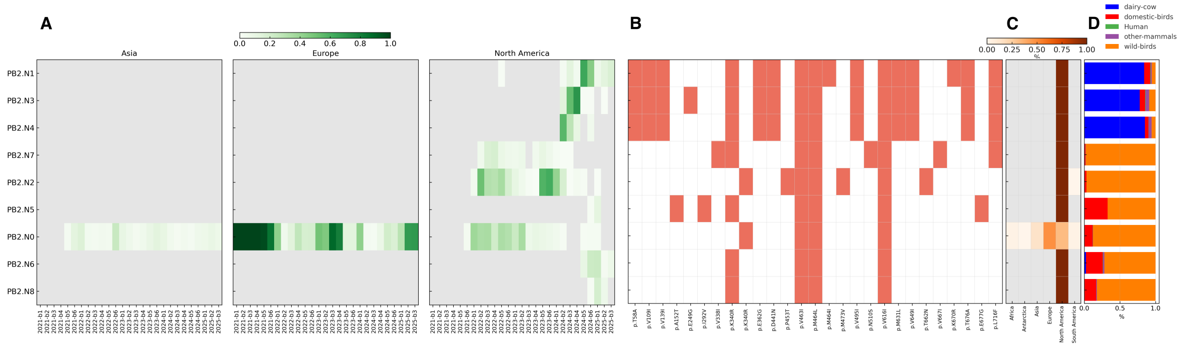


**Figure S7**: **Polymerase basic 2 (PB2): temporal and geographic distribution of groups.** (**A**) Temporal circulation of PB2 groups, shown as the fraction of sequences per bimester. (**B**) Heatmap of non-synonymous substitutions reveals two clusters: one linked to the B3.13 cattle genotype (PB2.N1, PB2.N3, PB2.N4) and one early cluster (PB2.N0). (**C**) Continental prevalence shows strong North American restriction, with PB2.N0 as the only globally distributed group. (**D**) Host distribution indicates that all B3.13-related PB2 groups share M631L, enhancing polymerase activity in mammals, and additional uncharacterized changes (T58A, V109I, V139I, E362G, D441N, V495I, V649I, T676A) that may influence replication efficiency.

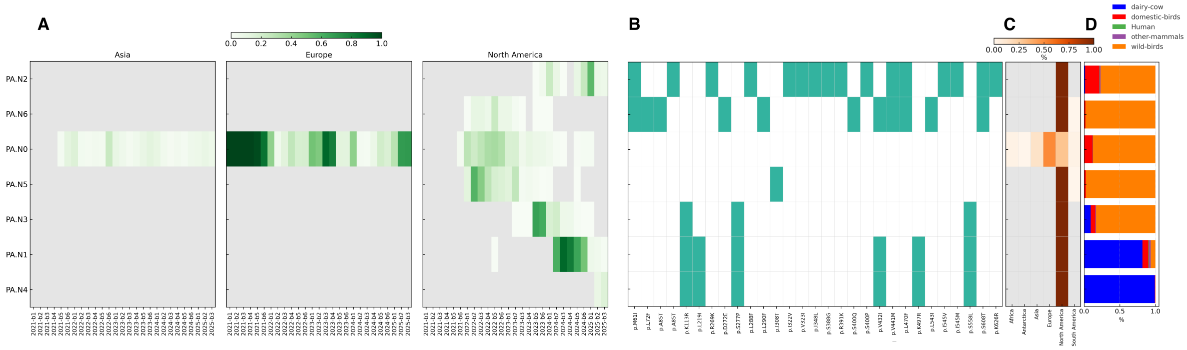


**Figure S8**: **Polymerase acidic (PA): temporal and geographic distribution of groups.** (**A**) Temporal circulation of PA groups, expressed as the fraction of sequences per bimester. (**B**) Heatmap of non-synonymous substitutions identifies four clusters: PA.N0 and PA.N5 (reference-like); PA.N1, PA.N3, and PA.N4 (B3.13-associated); and two divergent groups (PA.N2, PA.N6) restricted to North America. (**C**) Continental prevalence shows strong North American specificity for most groups. (**D**) Host association indicates that PA.N1 and PA.N4, linked to dairy cattle, carry K497R, which acts synergistically with PB2-M631L to enhance polymerase function in mammalian cells. PA.N3, found in early B3.13 and related genotypes, also carries this substitution.

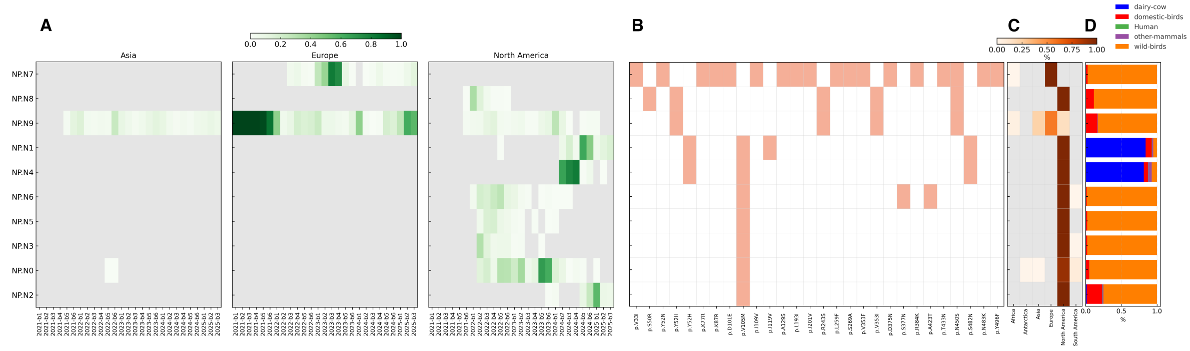


**Figure S9**: **Nucleoprotein (NP): temporal and geographic distribution of groups.** (**A**) Temporal circulation of NP groups, expressed as the fraction of sequences per bimester. (**B**) Heatmap of characteristic substitutions showing three clusters: NP.N9 and NP.N8; NP.N7; and groups NP.N0 to NP.N6. The latter cluster is restricted to North America and defined by V105M, a known mammalian-adaptive change. (**C**) Continental prevalence shows North American dominance for most NP groups. (**D**) Host association bar plot indicates that NP.N1 and NP.N4, linked to the B3.13 dairy cattle epidemic, carry Y52H (which confers escape from mammalian restriction factors MxA and BTN3A3) and S482N, a North American–specific variant.

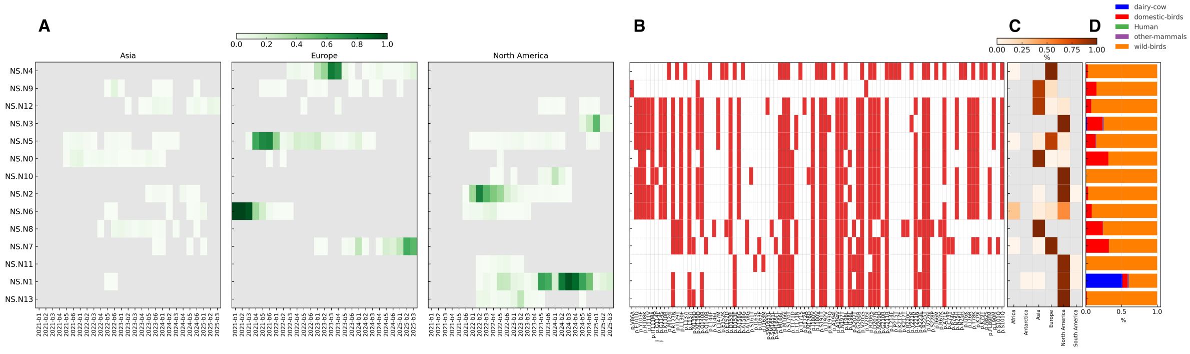


**Figure S10**: **Non-structural protein (NS): temporal and geographic distribution of groups.** (**A**) Temporal circulation of NS groups, expressed as the fraction of sequences per bimester. (**B**) Heatmap of non-synonymous substitutions identifies four clusters: an early group (NS.N12 and the related groups NS.N3, NS.N5, NS.N0, NS.N10, NS.N2, and NS.N6), a cluster including the two worldwide widespread groups NS.N1 and NS.N13, and the divergent NS.N4 and NS.N9, each restricted to Europe. (**C**) Continental prevalence shows broad intercontinental distribution, with limited geographic structuring compared to other segments. (**D**) Host association plot showing that NS variants infect a broad range of hosts without enrichment in any class.

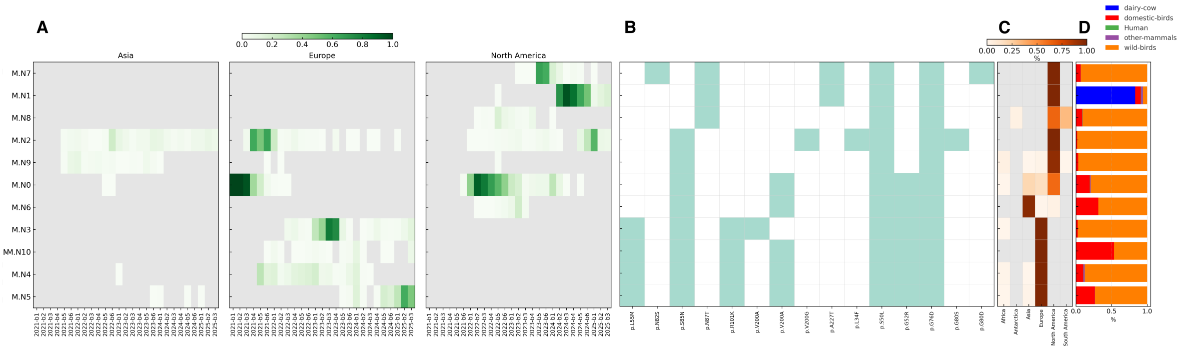


**Figure S11**: **Matrix (M): temporal and geographic distribution of groups.** (**A**) Temporal circulation of each M group, expressed as the fraction of sequences per bimester; only bimester–group combinations with available data are shown. (**B**) Heatmap of characteristic non-synonymous substitutions, with two main clusters: M.N2–M.N9, which share S85N and show widespread circulation, and M.N8–M.N1–M.N7, which share N87T, while M.N1 and M.N7 also carry A227T. (**C**) Continental prevalence highlights that M.N1, associated with the B3.13 dairy cattle genotypes, is restricted to North America. (**D**) Barplot of host associations showing proportional distribution of infections by host class. Although N87T and A227T remain uncharacterized in H5N1, their recurrence in North American viral grups suggests potential early steps toward local diversification.

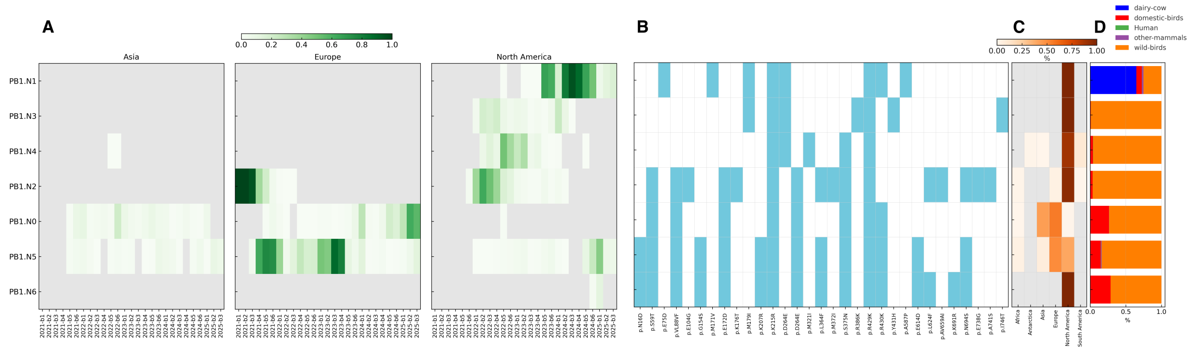


**Figure S12**: **Polymerase basic 1 (PB1): temporal and geographic distribution of groups.** (**A**) Temporal circulation of PB1 groups, shown as the fraction of sequences per bimester. (**B**) Heatmap of non-synonymous substitutions reveals two main clusters: PB1.N2–PB1.N6, globally distributed (except PB1.N6, specific to North America), and PB1.N1–PB1.N4, defined by D264E. (**C**) Continental prevalence highlights that PB1.N1, linked to the B3.13 cattle genotype, carries additional substitutions (E75D, M179I, Y431H, A587P). (**D**) Host distribution barplot. None of these PB1 substitutions are experimentally validated in H5N1, and polymerase studies do not identify them as known adaptation markers, suggesting they represent candidate sites for future functional work. Overall, PB1 mirrors PB2 and PA in its strong North American diversification.

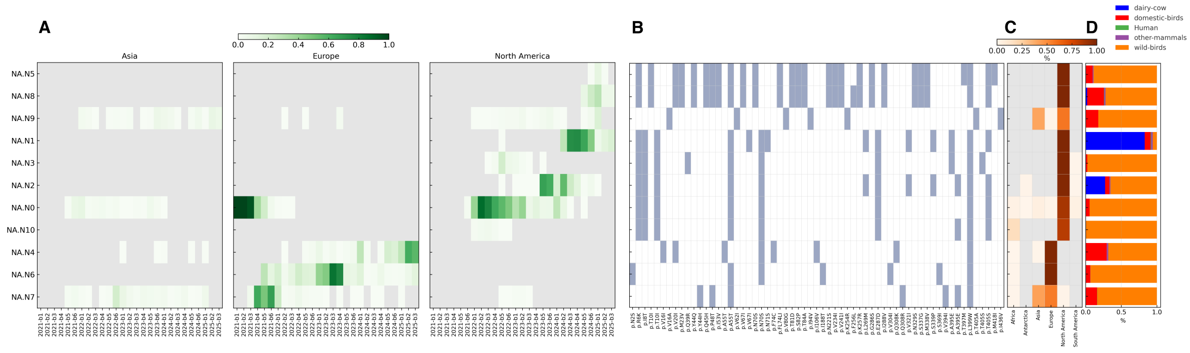


**Figure S13**: **Neuraminidase (NA): temporal and geographic distribution of groups.** (**A**) Temporal circulation of NA groups, expressed as the fraction of sequences per bimester; recently emerged clusters NA.N5 and NA.N8 appear only in the most recent time periods. (**B**) Heatmap of characteristic non-synonymous substitutions reveals four major macro-groups. NA.N5 and NA.N8 show an elevated number of amino acid changes, and NA.N8 is associated with the D1.1 genotype linked to the 2025 dairy cattle spillover. (**C**) Continental prevalence shows that North American groups are largely distinct from Eurasian and African clusters. NA.N9 and NA.N7 dominate in Asia, while NA.N6 and NA.N4 prevail in Europe. (**D**) Bar plot of host associations highlights dairy cattle–linked groups (NA.N1, NA.N2, NA.N8) carrying variants such as N71S and T405S in the 2SBS domain, potentially affecting receptor binding and host adaptation.

**Data S1.**

Metadata of 26,930 clade 2.3.4.4b viral isolates retrieved from the GISAID EpiFlu database. This table lists all isolates included in the study, along with associated metadata. Reported fields include: GISAID_ID (EPI_ISL identifier), Genotype (when available), Clade, Lineage, Host, Continent, Country, Region (geographic metadata, when available), Collection date, and Submission date. For each genomic segment (PB2, PB1, PA, HA, NP, NA, M, NS), the corresponding HaploCoV segment group is provided, as well as the combined haplogroup label (HC_comb).

**Data S2.**

Segment groups-defining amino acid changes. For each genomic segment (HA, M, NA, NP, NS, PA, PB1, PB2), the table lists all identified groups and their defining amino acid changes. Each segment–group combination is presented on a separate line.

**Data S3.**

Metadata of 25,779 isolates assigned to major viral groups included in downstream analyses. This table contains the subset of isolates classified into major viral groups used for subsequent analyses. In addition to the fields reported in Data S1, the following information is provided: WeekIso (year and ISO week of collection), hostClass (simplified host classification: wild birds, domestic birds, other mammals, dairy cows, or humans), and Label (group label following the nomenclature introduced in this study).

**Data S4.**

Major group features. For every group containing more than 20 isolates, the table reports key characteristics, including the segment-group assignments defined by HaploCoV for each of the eight genomic segments (PB2, PB1, PA, HA, NP, NA, M, and NS), the corresponding HC_comb label, the genotype, and the label assigned according to the rules described in the manuscript.

**Data S5.**

Domestic bird outbreaks. The table lists all groups for which at least 10% of the isolates and/or at least 10 total isolates were collected from domestic birds. For each group, the total number of isolates and the percentage originating from domestic birds (relative to the group total) are reported. Finally, we list all countries from which at least 10% of the isolates associated with spillover into domestic birds were detected. Countries are ranked according to the total number of isolates collected from domestic birds.

**Data S6.**

Non-synonymous substitutions. For each genomic segment (PB2, PB1, PA, HA, NA, NP, M, and

NS), the table reports the complete set of non-synonymous substitutions identified, along with their genomic coordinates, the reference sequence and amino acid, and the alternative sequence and amino acid. The presence or absence of each substitution across the segment groups defined by HaploCoV is indicated as 1 (present) or 0 (absent).
